# GSTU7 affects growth performance and acts as an antagonist of oxidative stress induced by methyl viologen

**DOI:** 10.1101/2020.06.09.142729

**Authors:** José Manuel Ugalde, Liliana Lamig, Ariel Herrera-Vásquez, Philippe Fuchs, Stefanie J. Müller-Schüssele, Andreas J. Meyer, Loreto Holuigue

## Abstract

Plant glutathione *S*-transferases (GSTs) are glutathione-dependent enzymes with versatile functions, mainly related to detoxification of electrophilic xenobiotics and peroxides. The Arabidopsis genome codes for 53 GSTs, divided into seven subclasses, however understanding of their precise functions is limited. A recent study showed that class II TGA transcription factors TGA2, TGA5 and TGA6 are essential for tolerance of UV-B-induced oxidative stress and that this tolerance is associated with an antioxidative function of cytosolic tau-class GSTUs. Specifically, TGA2 controls the expression of several GSTUs under UV-B light and constitutive expression of GSTU7 in the *tga256* triple mutant is sufficient to revert the UV-B-susceptible phenotype of *tga256*. To further study the function of GSTU7, we characterized its role in mitigation of oxidative damage caused by the herbicide methyl viologen (MV). Under non-stress conditions, *gstu7* null mutants were smaller than wild-type (WT) plants and delayed in the onset of the MV-induced antioxidative response, which led to accumulation of hydrogen peroxide and diminished seedling survival. Complementation of *gstu7* by constitutively expressed *GSTU7* rescued these phenotypes. Furthermore, live monitoring of the glutathione redox potential in intact cells with the fluorescent probe Grx1-roGFP2 revealed that *GSTU7* overexpression completely abolished the MV-induced oxidation of the cytosolic glutathione buffer compared to WT plants. GSTU7 was found to act as a glutathione peroxidase able to complement the lack of peroxidase-type GSTs in yeast. Together, these findings show that GSTU7 is crucial in the antioxidative response by limiting oxidative damage and thus protecting cells from oxidative stress.

## INTRODUCTION

In plants, animals and prokaryotes, glutathione *S*-transferases (GSTs) constitute a large family of enzymes with a wide range of cellular functions (Basantani and Srivastava, 2007). In plants, GSTs are mainly involved in the detoxification of exogenous xenobiotics and intracellular oxidized molecules, such as lipid peroxides, to alleviate chemical and oxidative damage (Sappl et al., 2009; Chronopoulou et al., 2017a) and anthocyanin transport to the vacuole (Alfenito et al., 1998; Mueller et al., 2000; Gomez et al., 2011). GST-mediated cellular detoxification is largely based on two enzymatic activities: the transferase and the peroxidase activities. As transferases, GSTs mediate the conjugation of the reduced tripeptide glutathione (GSH) with the electrophilic centre of different organic molecules (Dixon and Edwards, 2010; Labrou et al., 2015). The resulting conjugates are subsequently exported from the cytosol to the vacuole or the apoplast (Cummins et al., 2011). As peroxidases, some GSTs exhibit a GSH-dependent glutathione peroxidase (GPOX) activity which allows the reduction of lipid peroxides to alcohols (Edwards et al., 2000; Dixon et al., 2009). Plant GSTs are also involved in other processes such as secondary metabolism and development (Roxas et al., 1997; Chen et al., 2007; Dixon et al., 2010). Expression of several *GSTs* is boosted under biotic and abiotic stresses, e.g. pathogen infection (Mauch and Dudler, 1993), high salt concentrations (Roxas et al., 2000), hypoxia (Moons, 2003), UV-B radiation (Loyall et al., 2000), exposure to safeners (DeRidder et al., 2002; Cummins et al., 2009), herbicides (Dixon and Edwards, 2010), oxylipins (Mueller et al., 2008; Stotz et al., 2013) or salicylic acid (SA) (Chen et al., 1996; Sappl et al., 2009). The common denominator of these stress conditions is that they all trigger the production of reactive oxygen species (ROS) and hence cause oxidative stress. The induction of *GSTs* across this extensive span of conditions and treatments has strongly reinforced the idea of their protective role against oxidative stress (Chen et al., 2012; Chronopoulou et al., 2017b).

In *Arabidopsis thaliana*, the *GST* gene family counts 53 members of which only a few are functionally characterized (Sylvestre-Gonon et al., 2019). Based on their protein sequence and function, the GSTs are divided into seven distinct subclasses (Supplemental Fig. S1) (Wagner et al., 2002). The subclasses phi (F) and tau (U) are plant-specific and constitute the largest classes with 14 and 28 members, respectively (Chronopoulou et al., 2017a). Tolerance to herbicides mediated by GSTs relies on the capacity of GSTUs and GSTFs to conjugate electrophilic herbicides or their derivatives to GSH (Cummins et al., 1999; Edwards et al., 2000). Furthermore, studies on weeds indicate that herbicide resistance is mediated by GSTF isoforms and depends on preventing the accumulation of cytotoxic organic hydroperoxides caused by herbicide applications, rather than by the direct detoxification of the herbicide itself (Cummins et al., 1999).

The detoxification activities of GSTs strictly depend on GSH and thus are tightly interconnected with the cellular redox state (Dixon and Edwards, 2010). GSH is the most abundant low-molecular-weight thiol in the cell and is involved in a number of key cellular functions. Direct or indirect use of GSH as an electron donor for ROS scavenging leads to formation of glutathione disulfide (GSSG) (Noctor et al., 2012). Based on *in vivo* measurements with genetically encoded fluorescent probes, the GSH/GSSG ratio in subcellular compartments that contain a functional glutathione reductase has been calculated to be in the order of 50.000:1 (Schwarzländer et al., 2016). Stress situations and experimental systems like treatment with bacterial elicitors cause a pronounced oxidative shift (Nietzel et al., 2019). The shift in the GSH/GSSG ratio is a key feature of stress responses and is considered to be important for downstream signalling pathways and developmental programmes (Meyer, 2008).

Methyl viologen (MV) is a broad-spectrum herbicide that acts in chloroplasts by redirecting electrons originating from photosystem I (PSI) to molecular oxygen (O_2_), which generates superoxide (O_2_^⋅−^) (Mano et al., 2001; Scarpeci et al., 2008). The enhanced production of O_2_^⋅−^ has made MV a model for photooxidative stress in plants (Laporte et al., 2012). Similarly, in non-photosynthetic organisms, MV redirects electrons from complex I of the mitochondrial electron transport chain to O_2_, which also generates O_2_^⋅−^ (Cochemé and Murphy, 2008). Increased ROS production leads to lipid peroxidation, which is considered to be the main cause of MV toxicity in plant and mammalian cells (Bus et al., 1976; Misra and Gorsky, 1981; Liu et al., 2009).

Studies in plants have already demonstrated that GSTs contribute to MV tolerance. For instance, heterologous overexpression of cotton *GST-cr1* in tobacco (Yu et al., 2003), a *Suaeda salsa GST* in rice (Zhao and Zhang, 2006), and rice *GSTU4* in Arabidopsis (Sharma et al., 2014), as well as homologous overexpression of *GSTU51* in poplar (Choi et al., 2013), and *GSTU19* in Arabidopsis (Xu et al., 2016), consistently let to an improved MV tolerance when compared to their wild-type (WT) controls. These studies all noticed a connection between the tolerance phenotype and both an increased GPOX activity and a reduced ROS production upon MV treatment. The molecular function of GSTs in this context, however, remains unknown.

Expression of *GSTU7* has been reported to be induced in plants exposed to a wide range of stress stimuli including the exposure to heavy metals (Becher et al., 2004), to oxylipins and phytoprostanes (Loeffler et al., 2005; Mueller et al., 2008; Stotz et al., 2013) and to SA (Blanco et al., 2009). SA also promotes the accumulation of GSTU7 protein in cell cultures (Sappl et al., 2004; Gruhler et al., 2005). For oxylipin treatment, expression of *GSTU7* has been shown to be controlled by class II TGA transcription factors (TGA2, 5 and 6) (Stotz et al., 2013). Independent findings further support a role of class II TGA transcription factors in controlling the expression of a group of genes with putative antioxidant functions in response to SA (Blanco et al., 2009) and UV-B stress (Herrera-Vásquez et al., 2020). In the latter work, *tga2 tga5 tga6* triple mutants (*tga256*) were more susceptible to oxidative stress caused by UV-B similar to the susceptibility of *tga256* to SA established earlier by Zhang et al., 2003. Interestingly, at least 12 GSTUs including GSTU7 were induced by UV-B and shown to be controlled by the class II TGA transcription factors. The role of TGAs in expression of GSTU7 was further confirmed in ChIP assays showing binding of TGA2 to the palindromic TGA-box (TGACGTCA) in the *GSTU7* promoter region under stress conditions *in vivo* (Herrera-Vásquez et al., 2020). It is unclear though why GSTU7 would be beneficial in stress situations that activate class II TGAs.

The large number of GSTs in plants and widespread functional redundancy among them make it challenging to study the function of specific GSTs and therefore little is known about their individual roles in Arabidopsis. In this study, we report a role of GSTU7 in plant growth and characterize GSTU7 as a key enzyme mediating the tolerance of Arabidopsis to oxidative stress induced by MV and explore its importance in cellular redox homeostasis.

## RESULTS

### Expression of *GSTU7* is induced under different stress conditions

To determine *GSTU7* expression induced by different stress conditions, Arabidopsis WT seedlings were treated with either 0.5 μM SA, UV-B radiation or 50 μM MV and *GSTU7* transcript levels evaluated at different time points (Fig. 1, A-C). SA caused a six-fold induction of *GSTU7* expression within 30 min, leading to a peak of seven-fold expression after 2 h compared to non-treated controls (Fig. 1A). Between 8 to 24 h after the start of the SA incubation, *GSTU7* expression was again close to expression in control plants. UV-B radiation (Fig. 1B) induced a 2.3-fold expression after 2.5 h and slightly increased to three-fold at 24 h. Treatment of plants with MV resulted in the most pronounced induction of *GSTU7* expression among the three treatments with a maximum of 25-fold after 24 h (Fig. 1C).

**Figure 1.**
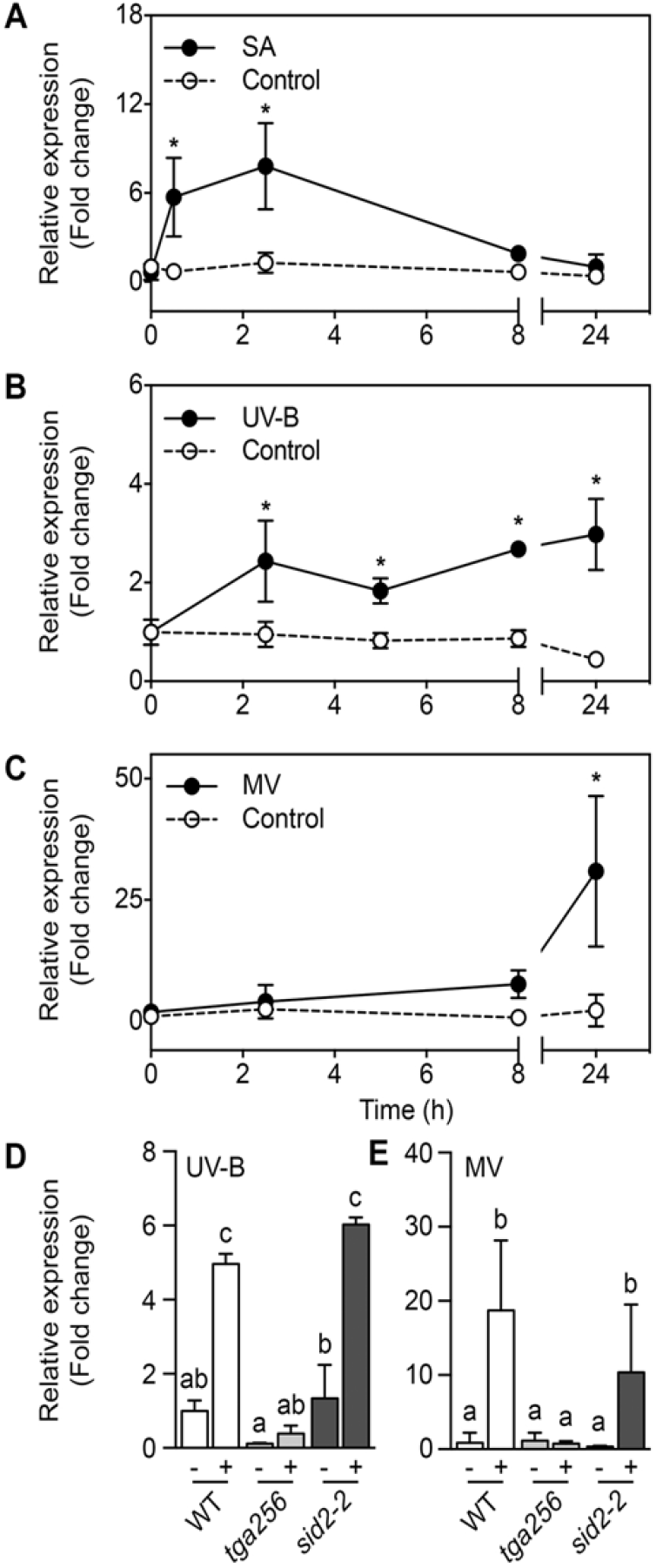
GSTU7 gene expression is induced by salicylic acid (SA), UV-B radiation (UV-B) and methyl viologen (MV) treatments. A-C, mRNA accumulation of *GSTU7* was determined by qPCR in 2-week-old WT Arabidopsis seedlings treated with 0.5 mM SA in MS medium (A), UV-B radiation (B) or 50 μM MV in MS medium (C) (all shown by closed circles). For control treatments (open circles), seedlings were floated on MS medium (A, C) or irradiated with UV-B under a glass filter (B). D-E, *GSTU7* transcript levels were measured by qPCR in WT, *tga256* and *sid2-2* seedlings after 5 h of UV-B radiation (D) or after 24 h of incubation with 50 μM MV (E). Gene expression is expressed as fold change relative to a subunit of *CLATHRIN ADAPTOR COMPLEX* (for SA and MV) or to *YLS8* (for UV-B) and normalized to the expression at 0 h (A-C) or to the expression in untreated WT seedlings (D-E). Means ± SD (A-E) were calculated from three independent biological replicates (2-3 seedlings pooled per replicate). The obtained data were tested for statistically significant differences by ANOVA with Fisher’s LSD test. * indicates differences compared to the expression at 0 h (*P*<0.05). Different letters indicate statistically different groups (*P*<0.05).

Recently, UV-B-induced expression of 12 GSTs including GSTU7 was found to be controlled by the class II TGA transcription factors TGA2, TGA5 and TGA6 (Herrera-Vásquez et al., 2020). To test whether the induction of GSTU7 in response to UV-B and MV (Fig. 1A) is dependent on class II TGAs or SA, UV-B- and MV-induced changes in *GSTU7* transcript levels were compared between WT, the *tga256* triple mutant and the SA-deficient mutant *sid2-2*. In contrast to the response in WT plants, induction of *GSTU7* expression was completely abolished in the *tga256* triple mutant (Fig. 1, D and E). In *sid2-2* mutant, however, expression of *GSTU7* was still induced by treatments with UV-B and MV, similarly to induction in WT plants (Fig. 1, D and E). Altogether, the experiments show that UV-B and MV stress-induced expression of *GSTU7* is controlled by class II TGAs but not through a SA-dependent pathway.

### Deletion of GSTU7 causes growth retardation

To further characterize the role of GSTU7, an insertional mutant, *gstu7,* was isolated and the T-DNA confirmed by sequencing to be inserted between A and T of the ATG start codon (Fig. 2A). Absence of detectable transcript confirmed *gstu7* as a null mutant (Fig. 2B). For complementation, a *UBQ10_pro_:GSTU7-V5* construct was cloned and transformed into *gstu7* plants. Two independent lines, *gstu7/GSTU7-V5* #2 and *gstu7/GSTU7-V5* #5 were selected from a larger pool of 16 transformants based on high expression levels and protein abundance. As expected, treatment of WT plants with 50 μM MV caused increased *GSTU7* expression, while in both complemented lines, *GSTU7* transcripts were detected at higher levels than in the WT, without pronounced changes after MV treatment (Fig. 2B). Constitutive expression of *GSTU7-V5* was confirmed by immunodetection of the V5 tag in both lines, albeit with much higher protein abundance in line #2 than in line #5 (Fig. 2C).

**Figure 2.**
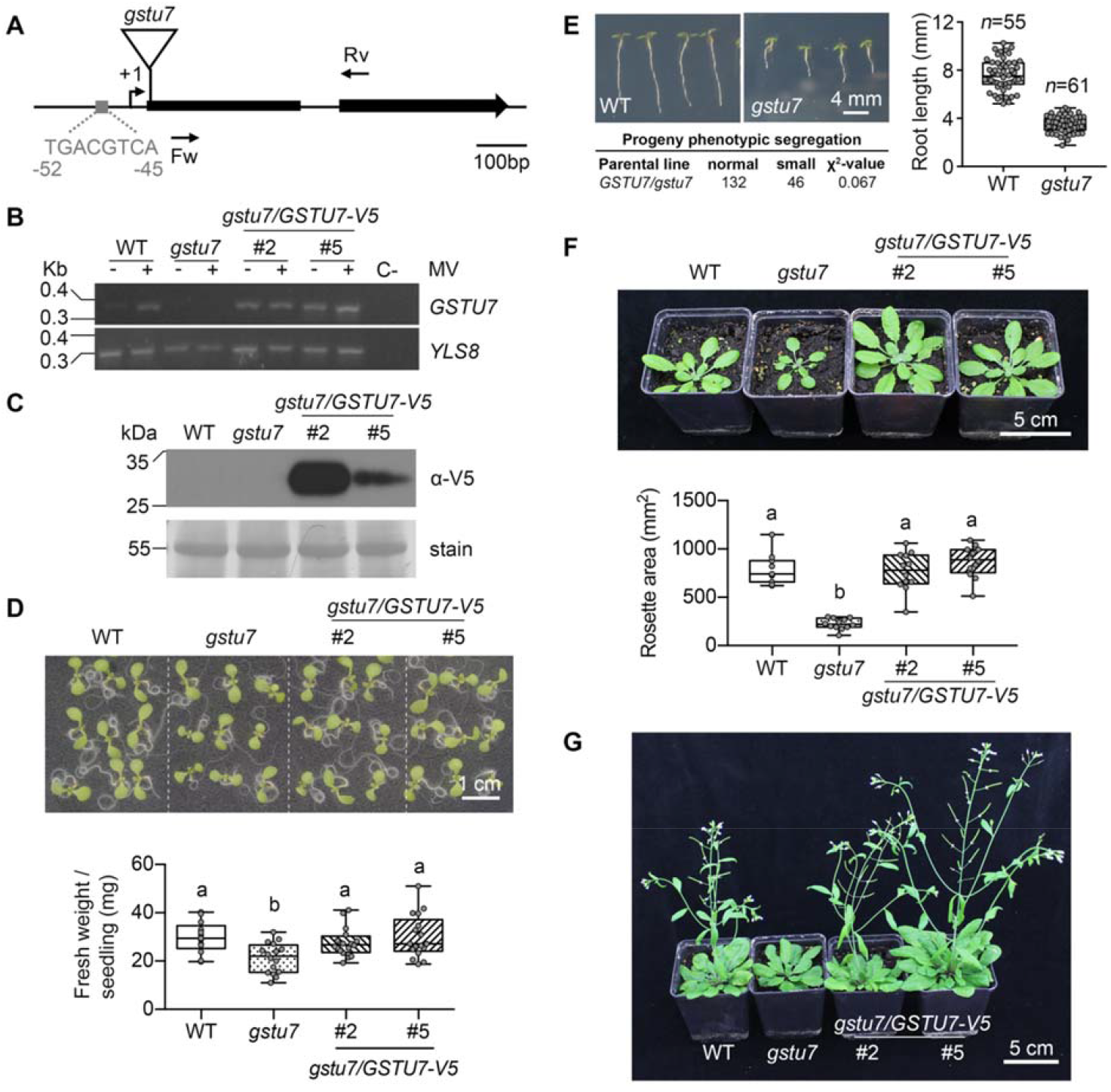
Characterization of the Arabidopsis *gstu7* mutant and the *gstu7/GSTU7-V5* complemented lines. A, Scheme of the *GSTU7* gene indicating the site of T-DNA insertion in the *gstu7* mutant line (SALK_086642). The TGA-box (grey) is around 50 bps upstream of the transcription start site (+1). Forward (Fw) and reverse (Rv) primer binding sites used for mRNA quantification are indicated. B, Quantification of *GSTU7* transcripts in WT, *gstu7* and two *GSTU7* overexpression lines. A fragment of the *YLS8* housekeeping gene was amplified as a control. C, Immunoblot analysis of GSTU7 using an antibody against the V5 tag of the complement. 20 μg protein were loaded per lane. To verify even loading, the membrane was stained with Coomassie (lower panel). D, Growth phenotype of seedlings. Photographs (upper panel) and fresh weight of whole seedlings (lower panel) one week after germination. Box plots show the mean of 13 to 17 pools of 4 to 6 seedlings, whiskers represent the min and max values. E, Segregation of *gstu7*-linked short root phenotype. WT and *gstu7* seedlings were grown vertically on plates and analysed for their root length on day 5 after stratification. Segregation of a backcross resulted in a short root:normal WT ratio of 1:3. Box plots show the mean of either 55 or 61 replicates, whiskers represent the min and max values. F, Rosette phenotypes of soil grown plants four weeks after germination. Box plots showing the mean of 12 replicates, whiskers represent the min and max values. G, Appearance of the mutant and complemented lines seven weeks after germination. Statistical analyses for panels D, E and F were performed using ANOVA with Fisher’s LSD test. Different letters indicate statistically different groups (*P*<0.05).

Homozygous *gstu7* mutants developed a dwarf phenotype with lower fresh weight compared to WT seedlings (Fig. 2D), shorter roots (Fig. 2E), smaller rosettes (Fig. 2F), and delayed bolting (Fig. 2G). Complementing the mutant with *GSTU7-V5* restored *gstu7* to a WT phenotype at all analysed developmental stages. To confirm that the smaller phenotype of *gstu7* was due to a single T-DNA insertion, *gstu7* was backcrossed to a WT plant. The F2 progeny of this cross segregated with a 3:1 ratio for WT:short root mutant phenotypes (χ^2^=0.067; *P*=0.79) (Fig. 2E).

### GSTU7 mitigates oxidative stress induced by methyl viologen

The main function of GSTs is the detoxification of exogenously applied xenobiotics and endogenously generated organic peroxides, such as lipid peroxides (Sappl et al., 2009; Chronopoulou et al., 2017a). To further test whether GSTU7 is involved in developing plant tolerance to oxidative stress, seed germination of WT, *gstu7* and the two complemented lines #2 and #5 was investigated in MS plates with or without 0.1 μM MV (Fig. 3A). Seedling tolerance to MV was determined as survival rates, i.e. the percentage of seedlings with green tissue out of all germinated seedlings (Laporte et al., 2012). *gstu7* showed a survival rate of 29% compared to WT seedlings of which 62% survived (Fig. 3A). Constitutive overexpression of GSTU7 cured the apparent increased susceptibility of *gstu7* towards MV and led to a survival rate of about 90%, which is significantly higher than in WT plants (*P*=0.0003).

**Figure 3.**
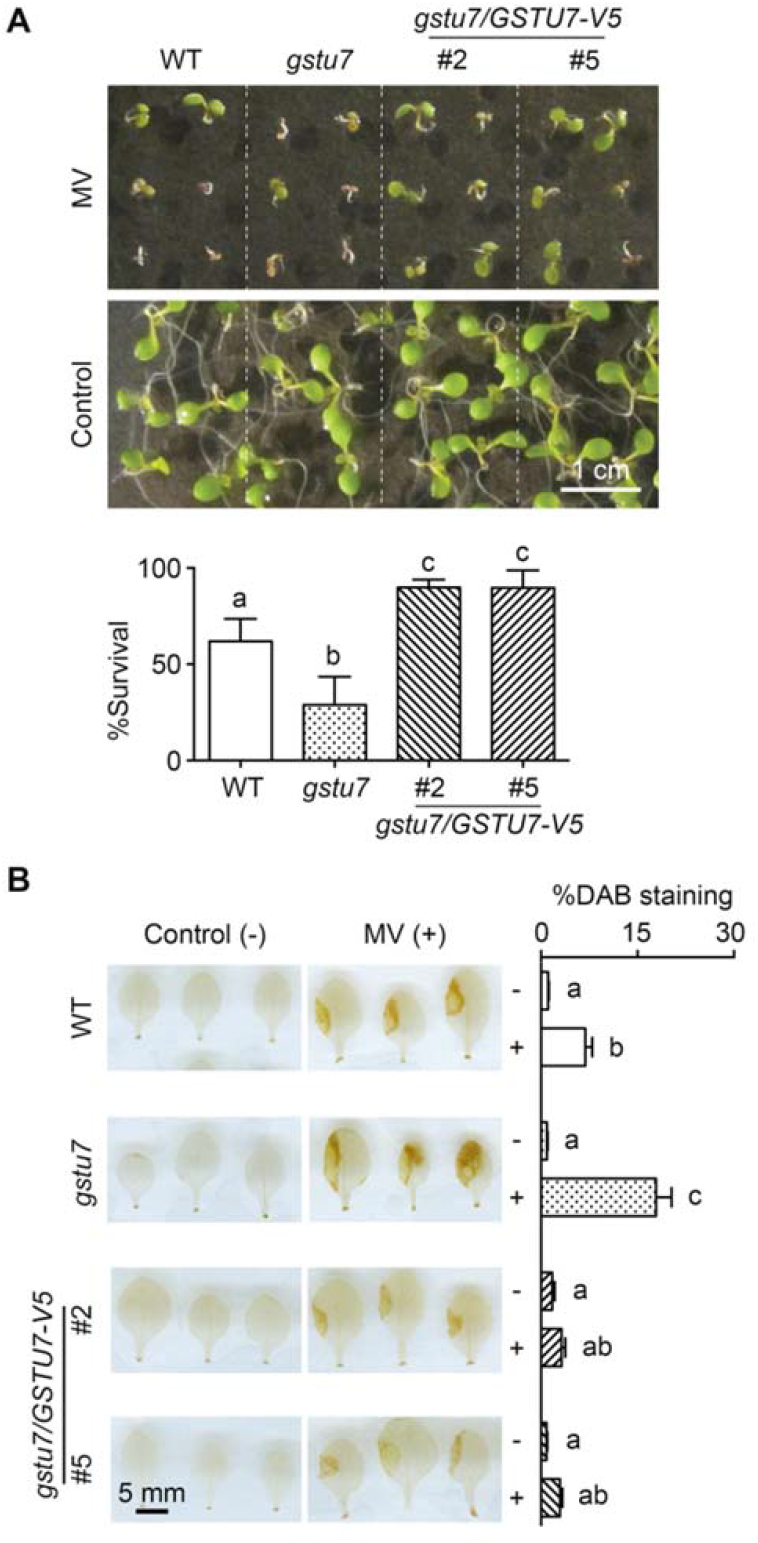
GSTU7 counteracts oxidative stress induced by MV. A, Phenotype of seedlings from WT, *gstu7* and *gstu7/GSTU7-V5* complemented lines (#5 and #2) grown on plates with or without 0.1 μM MV. Seedling tolerance to MV treatment was measured after one week as the survival rate, which is the percentage of plants with any green cotyledon over the total number of germinated seeds (*n*=3, means +SD). B, Accumulation of H_2_O_2_ in primary leaves. Leaves of plants from different genotypes were exposed to MV through placement of a drop (2 μL) of 30 μM MV on the leave surface and kept under constant light for 24 h. Subsequently, H_2_O_2_ accumulation was evaluated by staining with DAB and expression as the fraction of intensely stained leaf area relative to the whole leave area (% DAB staining) (*n*=9, means + SD). Statistical analyses were performed by ANOVA with Fisher’s LSD test. Different letters indicate statistically different groups (*P*<0.05).

To further assess whether the increased susceptibility of *gstu7* mutants to MV correlates with a higher production of ROS, primary leaves were treated with a 2 μL drop of 30 μM MV, incubated under constant light conditions for 24 h and subsequently stained with 3,3’- diaminobenzidine (DAB) for hydrogen peroxide (H_2_O_2_) detection (Fig. 3B). For a semi-quantitative assessment, the leaf area stained with DAB was determined as the 3% more intense pixels relative to the area of the whole leaf (Fig. 3B). After MV treatment, *gstu7* seedlings had a three-fold larger area of intense staining than WT seedlings, which indicates the presence of increased amounts of H_2_O_2_. In contrast, *gstu7* seedlings overexpressing *GSTU7* after MV treatment showed even less staining than WT seedlings. In this case, the staining was not significantly different from staining in control seedlings not treated with MV (*P*<0.0001) (Fig. 3B).

A similar stress-induced increase in DAB staining was recently reported for *tga256* after exposure to UV-B light (Herrera-Vásquez et al., 2020). Because the respective TGA transcriptions factors control the expression of *GSTU7* (Fig. 1E), we tested whether *tga256* is also more sensitive to MV and whether this sensitivity is mitigated by overexpression of GSTU7. Consistent with the hypothesis, *tga256* mutants showed a survival rate of only about 4% compared to about 45% for the WT on plates containing 0.1 μM MV (Fig. 4A). Constitutive overexpression of GSTU7 in *tga256* increased the survival to WT level plants and thus rescued the hypersensitivity of *tga256* to MV. Taken together, these results strongly suggest that GSTU7 plays a central role in mitigating the oxidative stress response triggered by MV.

**Figure 4.**
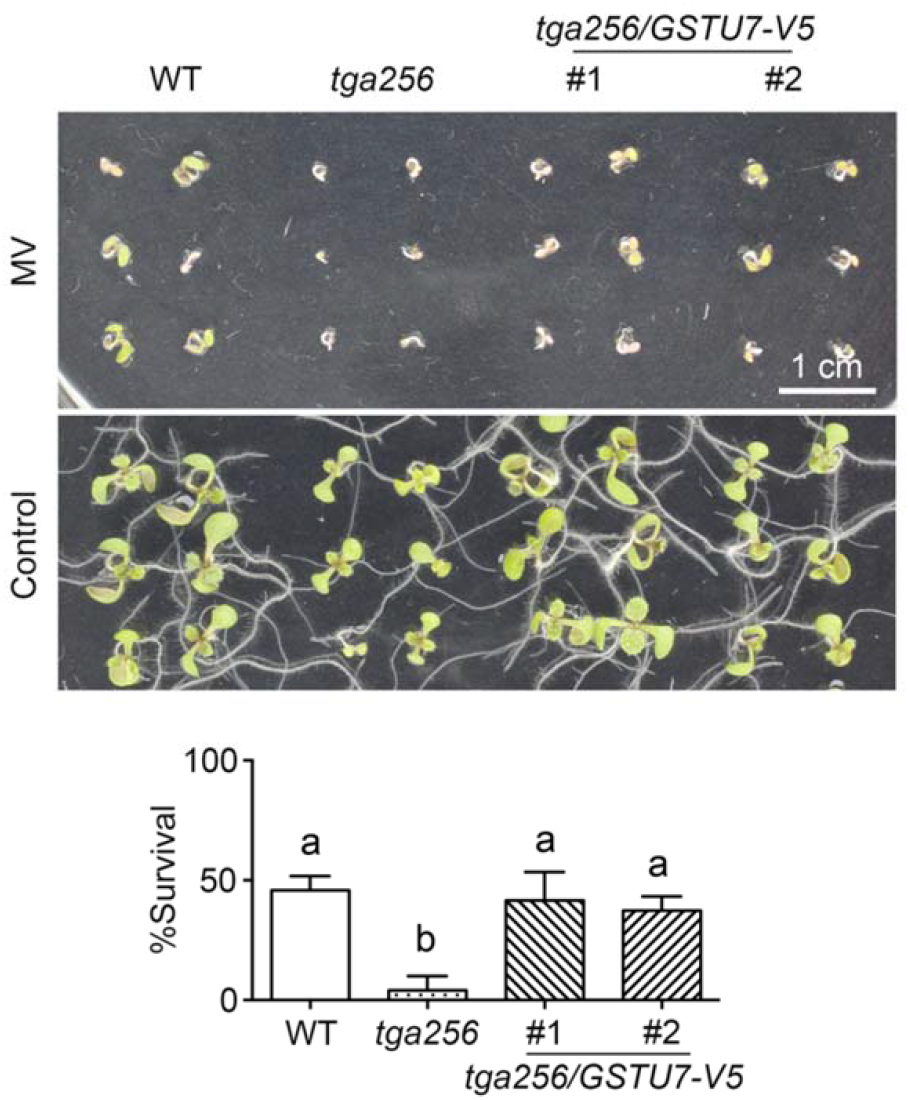
GSTU7 rescues *tga256* triple mutants from hypersensitivity to MV. Seeds of WT, *tga256* and *tga256/GSTU7-V5* complemented lines (#1 and #2) were sown on MS plates supplemented with or without 0.1 μM MV. Seedling tolerance to MV was measured after one week as the survival rates. Data indicate the mean values + SD from three independent experiments, where for each experiment 30 germinated plants were counted. Statistical analysis was performed using by ANOVA with Fisher’s LSD test. Different letters indicate different groups (*P*<0.05).

### GSTU7 is located in the cytosol and has glutathione peroxidase activity

In Arabidopsis, most GSTs are predicted to be localized in the cytosol (Supplemental Fig. S1). GSTU7 has been found in the Arabidopsis cytosolic proteome (Ito et al., 2011) and a GSTU7-GFP fusion has only been mentioned to be localized in the cytosol (Dixon et al., 2009). To further confirm these findings, we transiently expressed a GSTU7-GFP fusion protein in tobacco leaves and visualized the localisation on a confocal microscope (Fig. 5A). All images consistently revealed cytosolic localization of the fusion protein. In all observations, the nuclear region was marked by a bright ring surrounding a dim region with low fluorescence indicating that the fusion protein is barred from the nucleoplasm.

**Figure 5.**
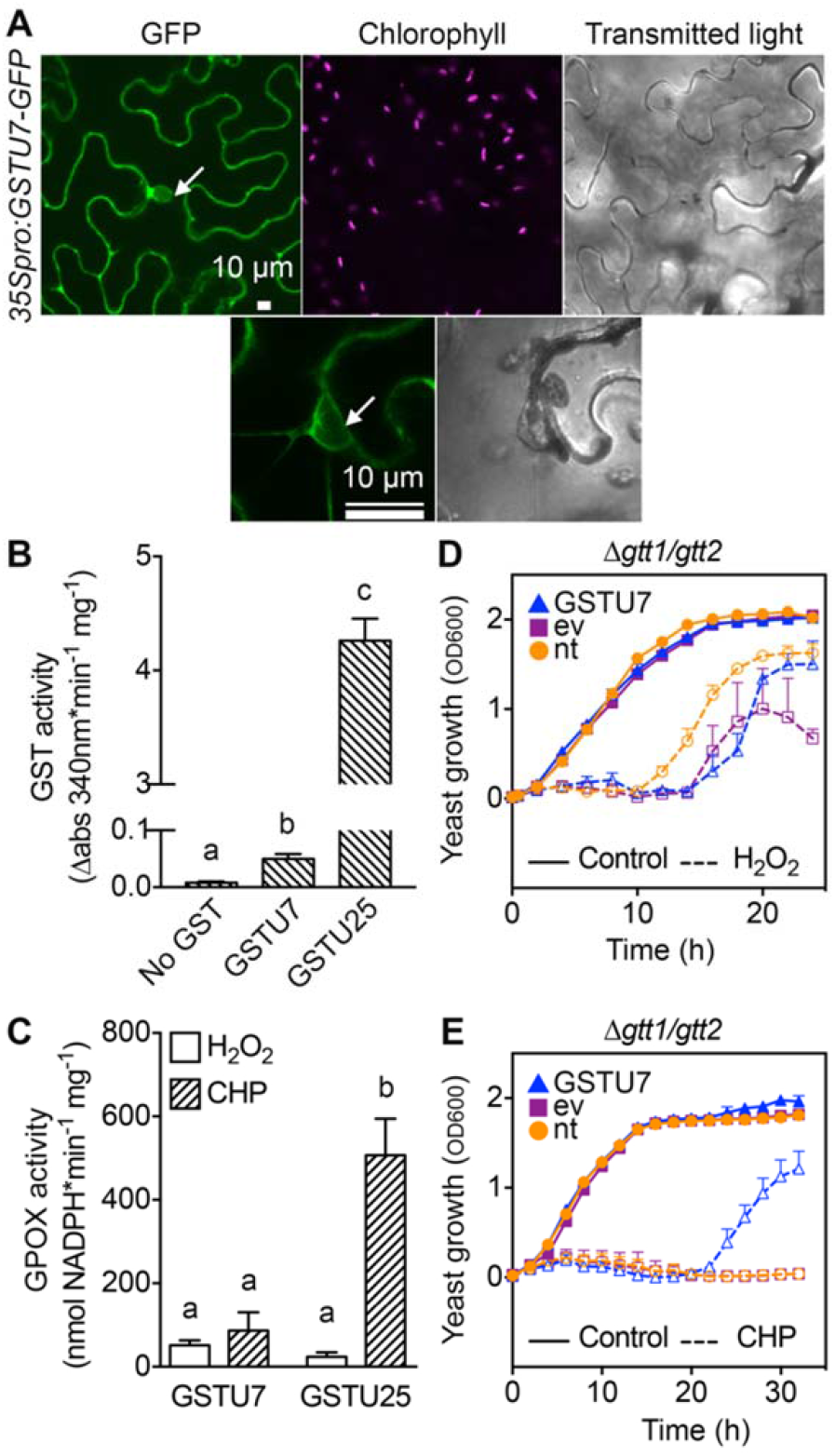
GSTU7 is located in the cytosol and has glutathione peroxidase (GPOX) activity. A, Tobacco leaves transiently expressing *35S_pro_:GSTU7-GFP* imaged 48 h after infiltration. Arrow heads indicate the nucleus. Bars = 10 μm. B, GST activity of recombinant GSTU7 or GSTU25 (mean + SD of *n*=21 for GSTU7 and *n*=3 for GSTU25). C, GPOX activity of GSTU7 and GSTU25 measured as consumption of NADPH in a coupled assay with glutathione reductase reducing the produced GSSG (mean value + SD of *n*=4–9). Statistical analysis in panels B and C was performed using by ANOVA with Fisher’s LSD test. Different letters indicate statistically different groups (*P*<0.05). D-E, Heterologous expression of GSTU7 in a Δ*gtt1/gtt2* yeast strain deficient in two GSTs. Yeast cells were incubated in liquid YPD supplemented with 1.5 mM H_2_O_2_ (D) or 0.09 mM CHP (E) (open symbols; dashed lines). Growth of non-treated controls is marked with closed symbols and solid lines. nt: non-transformed; ev: empty vector control.

Because of the pronounced mitigation of oxidative stress in GSTU7 overexpressing seedlings (Fig. 3B), the enzymatic activity of GSTU7 was further evaluated *in vitro* with the recombinant protein and 1-chloro-2,4-dinitrobenzene (CDNB) as an electrophilic substrate for GST activity and with either H_2_O_2_ or cumene hydroperoxide (CHP) as substrates for a putative GPOX activity (Fig. 5, B and C). GSTU25, which has been reported to be the GST with the highest GPOX and GST activities in Arabidopsis (Dixon et al., 2009), was used as a reference. GSTU7 showed a minor GST activity and was outperformed 97-fold by GSTU25 (Fig. 5B). Both GSTU7 and GSTU25 were capable of reducing H_2_O_2_ with activities between 51.3 and 23.5 nmol NADPH min^−1^ mg^−1^ (Fig. 5C). Activity of GSTU7 towards CHP was 87.3 nmol min^−1^ mg^−1^ and thus in the same order of magnitude as towards H_2_O_2_. In contrast, GSTU25 revealed a six-fold higher turnover of CHP compared to H_2_O_2_ (Fig. 5C).

To validate that the GPOX activity of GSTU7 observed *in vitro* has biological relevance *in vivo*, we tested whether GSTU7 can complement a yeast mutant deficient in the respective activity. The Δ*gtt1/gtt2* mutant in *Saccharomyces cerevisiae* lacks two key GSTs and has been reported to be sensitive to H_2_O_2_ and organic peroxides (Choi et al., 1998; Collinson and Grant, 2003) (Fig. 5D and E). This yeast strain was transformed with either *GSTU7* or an empty vector control. In the absence of exogenous oxidants, transformed and non-transformed cultures showed the same logarithmic growth curves. Supplementing YPD with either 1.5 mM H_2_O_2_ (Fig. 5D) or 0.09 mM CHP (Fig. 5E) abolished growth of the cultures almost completely. In medium supplemented with H_2_O_2_, growth started only after a lag phase of 10 h. Cells expressing GSTU7 showed an even longer lag phase of 14 h but reached the same density after 24 h (Fig. 5D). Incubation with CHP completely abolished growth of non-transformed cultures while Δ*gtt1/gtt2* cells expressing GSTU7 were able to survive and regain growth after 20 h (Fig. 5E). This supports the hypothesis that GSTU7 possesses GPOX activity *in vivo*, especially towards CHP.

Because selected glutathione peroxidase (GPX) proteins have been shown to act as protein thiol oxidases and thus can be exploited as H_2_O_2_ sensors (Delaunay et al., 2002; Gutscher et al., 2009), we also tested whether GSTU7 also is capable of transferring the primary oxidation of the catalytic cysteine resulting from peroxide detoxification to roGFP2 as a target protein. In contrast to bovine GPX as a positive control, GSTU7 was not able to mediate a H_2_O_2_-dependent oxidation of roGFP2 (Supplemental Fig. S2).

### GSTU7 protects cytosolic glutathione from MV-induced oxidation

Massive accumulation of ROS induced by MV treatment should also lead to a gradual oxidation of the glutathione redox buffer in the cell. With Grx1-roGFP2 as a fully dynamic probe for the local *E*_GSH_, the impact of MV and the role of GSTU7 on the oxidative load in the cytosol can be monitored in living cells over extended time periods. To test whether GSTU7 protects the cell from increased and ultimately deleterious oxidation, we monitored the cytosolic *E*_GSH_ with Grx1-roGFP2 *in vivo*, using WT, *gstu7* and *gstu7*/*GSTU7-V5* (#5 line) plants as genetic background. Stable homozygous roGFP2 reporter lines were selected and confirmed to show sufficiently high fluorescence intensities and typical spectral properties of the sensor with two excitations peaks and ratiometric properties using either a confocal microscope or a plate reader (Supplemental Fig. S3, A-C).

Changes in the roGFP2 fluorescence ratio triggered by MV treatments were measured in leaf discs of WT, the *gstu7* mutant and one of the complemented lines all expressing the Grx1-roGFP2 sensor (Fig. 6, A-C). Considering that manipulation of the leaf discs itself can trigger a stress response resulting in an altered redox state (Rosenwasser et al., 2010), leaf samples were initially incubated with 10 mM dithiothreitol (DTT) to attain complete and uniform reduction of the sensor. With this precaution, all measurements started with the sensor at the same level of reduction, serving as the reference point for later ratio normalization. After pre-reduction of the sensor, DTT was removed and leaf discs were incubated either with 100 μM MV or with water as control. WT plants treated with MV showed a gradual increase in the fluorescence ratio over time indicating gradual oxidation of the glutathione pool (Fig. 6A, Supplemental Fig. S4). This increase in the fluorescence ratio was more pronounced in *gstu7* plants (Fig. 6B, Supplemental Fig. S4). In contrast, leaves expressing GSTU7 showed no MV-induced oxidation and Grx1-roGFP2 remained almost completely reduced throughout the entire time course of 15 h (Fig. 6C, Supplemental Fig. S4).

**Figure 6.**
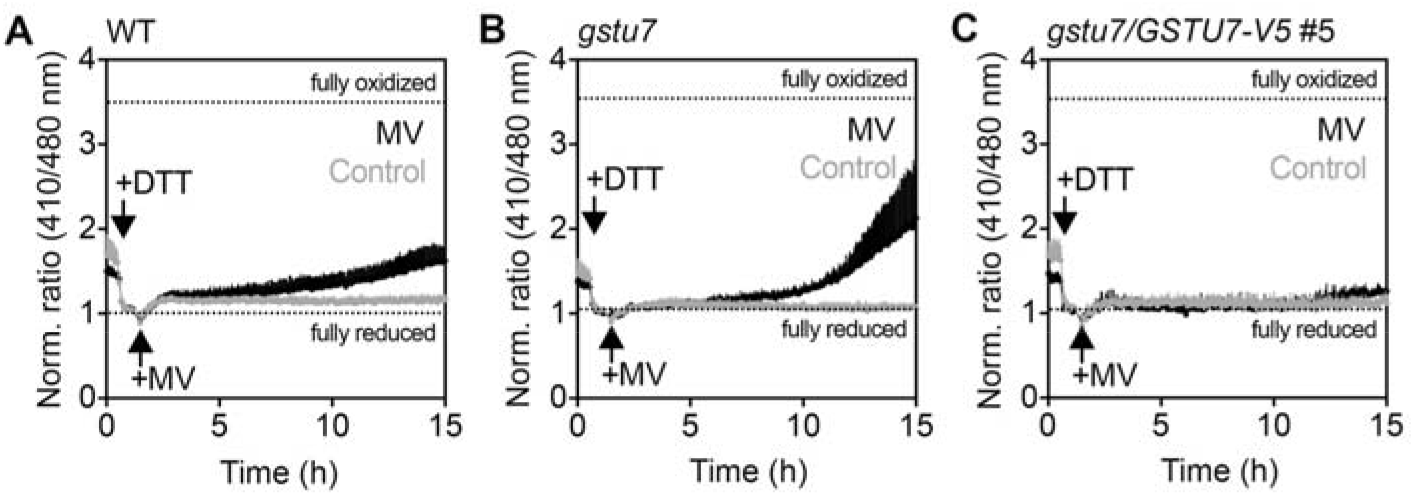
GSTU7 prevents MV-induced oxidation of cytosolic glutathione. A-C, Grx1-roGFP2 was expressed in Arabidopsis WT, *gstu7* and *gstu7/GSTU7-V5* #5 backgrounds. Leaf discs from stably transformed plants were placed in a 96-well plate and treated with 10 mM DTT to fully reduce the sensor (first black arrow). Subsequently, DTT was removed and the samples were treated with either 100 μM MV or water as control (second black arrow). The redox state of the sensor was followed over time as the ratio between the fluorescence excited at 410 and 480 nm. Ratio values were normalized to the fully reduced sensor after addition of DTT. The dotted lines indicate the ratios for fully reduced and fully oxidised roGFP2 measured at the end of the time course. All data indicate the mean fluorescence + SD of six samples from independent plants.

## DISCUSSION

It is well established that *GSTU7* is strongly induced under different abiotic stress situations and that many GSTs are induced by SA and other phytohormones (Marrs, 1996; Sylvestre-Gonon et al., 2019). Our results confirm that SA induces the expression of *GSTU7* within a few hours of application. In addition, there is also a medium to long-term response resulting in a very pronounced induction of *GSTU7* expression 5 hours after irradiation with UV-B or 24 hours after application of MV (Fig. 1D). Both transcriptional responses are mediated by class II TGAs but are independent of the endogenous production of SA (Supplemental Fig. S2). This indicates that *GSTU7* expression is controlled by at least two independent signalling pathways. A common response to stress situations is the formation of ROS resulting in secondary responses like lipid peroxidation generally summarized as oxidative stress (Waszczak et al., 2018; Smirnoff and Arnaud, 2019). UV-B triggers an accumulation of H_2_O_2_ and the finding that GSTU7 can complement the UV-B-sensitive phenotype of *tga256* led to the hypothesis that GSTU7 has an antioxidative function (Herrera-Vásquez et al., 2020). Treatment with MV is a classical means of inducing oxidative stress in green tissues because MV transfers electrons from photosystem I to molecular O_2_ to form superoxide (O_2_^⋅−^) (Mano et al., 2001). Increased susceptibility of *gstu7* null mutant plants to MV and the more intense MV-induced DAB staining was completely reverted by overexpression of *GSTU7* (Fig. 3). Although it cannot be fully excluded that DAB staining is also responsive to increased level of phenolic compounds (Veljovic-Jovanovic et al., 2002), all evidences presented here are consistent with the hypothesis that more intense DAB staining is a reliable proxy for the increased production of H_2_O_2_. It cannot be distinguished though whether the increased production of H_2_O_2_ is a primary or a secondary effect of MV application.

Deletion of GSTU7 resulted in a dwarf phenotype, which was consistently smaller than WT plants throughout its life cycle (Fig. 2, D-G). Delayed bolting of *gstu7* is similar to phenotypes of *gstf2* knockdown mutants and *gstu17* null-mutants (Gong et al., 2005; Jiang et al., 2010; Chen et al., 2012). While loss of GSTU17 expression has no impact on the development of young seedlings under control conditions (Jiang et al., 2010), loss of GSTU7 appears to affect the plant in a more general form resulting in defects in all developmental stages. Conversely, overexpression of a *Brassica campestris GSTU* closely related to Arabidopsis GSTU24 in Arabidopsis resulted in bigger rosettes, longer primary roots and an increased number of lateral roots (Kao et al., 2016). At the molecular level, GSTU17 controls plant development by regulating far-red light signalling through interaction with phytochrome A (*phyA*) (Jiang et al., 2010). Recently, selective binding of natural products has been shown for GSTU19 and GSTF2 from Arabidopsis (Dixon and Edwards, 2018). Although recombinant GSTU7 was found to preferentially retain glutathionylated protoporphyrin, no binding of unique ligands was found after heterologous expression in tobacco leaves (Dixon and Edwards, 2018).

It is established that GSTUs and GSTFs are subject to complex transcriptional and post-translational regulation in response to infection, abiotic stress, and development (Gullner et al., 2018; Kumar and Trivedi, 2018; Sylvestre-Gonon et al., 2019). The list of abiotic stress factors affecting GST expression has recently expanded to UV-B, which induces the expression of 12 *GSTUs* including *GSTU7* via class II TGA transcription factors (Herrera-Vásquez et al., 2020). It is tempting to speculate about a function of GSTU7 in ROS metabolism and signalling because the balance of peroxidase activity in roots has shown that H_2_O_2_ is involved in the control of root growth (Tsukagoshi et al., 2010). The nature of the GSTU7 function indicated here for non-stress condition, however, remains completely unknown and deserves further investigation in the future.

Previous studies showed that GSTs mediate detoxification of several classes of herbicides (Timmerman, 1989; Prade et al., 1998) by conjugating GSH to the electrophilic centre of these chemical compounds and that plants overexpressing GSTs are more resistant to herbicides (Cummins et al., 1999; Cummins et al., 2013). Furthermore, plant tolerance to MV was increased by overexpressing thylakoid ascorbate peroxidase (Murgia et al., 2004), copper-zinc superoxide dismutases (Furusawa et al., 1984; Perl et al., 1993; Aono et al., 1995), as well as different Arabidopsis GSTs, like GSTU4 (Sharma et al., 2014) and GSTU19 (Xu et al., 2016). MV has no electrophilic site for conjugation with GSH and its elimination in plants is thought to take place primarily via photocatalytic degradation (Moctezuma et al., 1999) or through the action of the yeast *Lipomyces starkeyi* when MV enters the soil (Carr et al., 1985). In mammals and plants, MV toxicity is associated with the accumulation of lipidperoxides (Misra and Gorsky, 1981; Liu et al., 2009). Efficient elimination of MV-induced secondary cytotoxic compounds like lipidperoxides is therefore assumed to be a critical component of the cellular tolerance to the herbicide and could be a key function of GSTs in the defence response.

GSTs are mostly located in the cytosol, with only 3 family members predicted to be localized in plastids (Supplemental Fig. S1). We confirmed the cytosolic localization of GSTU7 for heterologous expression in tobacco leaves (Fig. 5A). In line with observations by Dixon et al. (2009) we found GSTU7 being excluded from the nucleus. But in the absence of any obvious nuclear exclusion signal it is not clear how this exclusion is mediated or whether it is a consequence of the GFP fusion, which results in a larger fusion protein of 53 kDa. Especially if GSTU7 would form homo- or heterodimeric protein complexes the molecular mass would be more than 100 kDa, which is well above the size exclusion of about ~40 kDa of nuclear pores for passive transport (Tamura and Hara-Nishimura, 2014).

Most GSTUs have been reported to have a GPOX activity with CPH as substrate (Dixon et al., 2009). This includes GSTU7, which, however, is among the family members with the lowest GPOX activity. Our data confirm this low activity also in comparison to the highly active GPOX GSTU25. Similarly, the about 100-fold lower GST activity towards CDNB as an electrophilic substrate for GSTU7 compared to GSTU25 largely confirms the relative activities measured by Dixon et al. (2009). As a different approach, we tested for thiol oxidase activity of GSTU7 with roGFP2 as the reporting target protein for oxidation. All these attempts with recombinant GSTU7, however, failed and thus do not provide a clear lead for further biochemical characterization of the enzymatic activity. This leaves us with the conundrum of an enzyme preventing cells from severe oxidation induced by MV or UV-B and a lack of biochemical information. A possible limitation of the biochemical *in vitro* assays may be that GSTs tend to form homo- or heterodimeric protein complexes (Cummins et al., 1997; Dixon et al., 1997; Gronwald and Plaisance, 1998), which may not occur with recombinant proteins. In this situation, we thought to get access to the function of GSTU7 by testing its activity *in vivo*. In a previous study that aimed at functionally characterizing the large family of plant GSTs for their role in fungicide detoxification, almost all Arabidopsis GSTs were expressed in a GST-deficient yeast strain (Δ*gtt1* Δ*gtt2* Δ*grx1* Δ*grx1* Δ*tef4*), which is hypersensitive to fungicides and CDNB (Krajewski et al., 2013). Complementation of this yeast strain with different GSTs, including GSTU7, resulted in increased tolerance towards fungicides. Here overexpression of GSTU7 in the GST-deficient yeast strain Δ*gtt1* Δ*gtt2* conferred at least partial tolerance to CHP, which supports the notion that GSTU7 may be involved in lipidperoxide detoxification. roGFP2 is extremely sensitive to even minor increases in glutathione disulfide (GSSG) during detoxification of peroxides with GSH as electron donor (Marty et al., 2009). The completely abolished MV-induced oxidation in GSTU7 overexpressor plants strongly argues against the hypothesis for a GPOX activity and rather suggests that GSTU7 acts earlier in the pathway between MV and H_2_O_2_ accumulation. The fact that GSTU7 also rescues *tga256* mutants from UV-B stress (Herrera-Vásquez et al., 2020) strongly argues for a direct role of GSTU7 in peroxide detoxification.

Overall, our results reveal the importance of Arabidopsis GSTU7 as a protective enzyme for cellular detoxification, development and the antioxidant response to herbicides, such as MV. Yet, further studies are needed to elucidate the mechanistic properties of GSTU7 during MV-induced oxidative stress. These findings open new research possibilities for breeding of more stress-tolerant plants, in which over-expression of *GSTs* may contribute to overcoming present and future challenging environmental conditions that would lead to formation of ROS in plants.

## MATERIALS AND METHODS

### Plant material and growth conditions

Arabidopsis (*Arabidopsis thaliana* [L.] Heynh.) wild-type (WT), *sid2-2* (Wildermuth et al., 2001), *tga256* (Zhang et al., 2003), *gstu7* (SALK_086642C, ABRC, characterized in this work), *tga256*/GSTU7-V5 lines #1 and #2 (Herrera-Vásquez et al., 2020) and *gstu7*/*GSTU7* lines #2 and #5 (obtained in this work) were in Columbia (Col-0) background. Seeds were surface sterilized with 70% ethanol and rinsed three times with sterile deionized water. Clean seeds were sown on plates with 0.5x Murashige and Skoog (MS) growth medium (Murashige and Skoog, 1962) (Phyto Technology Laboratories, Shawnee Mission, KS) supplemented with 10 g L^−1^ sucrose and 3 g L^−1^ phytagel (Sigma, Steinheim, Germany) or 8 g L^−1^ agar (Winkler, Santiago, Chile). Plates were incubated in a growth chamber under a long day regime (16 h light, 100 μmoles m^−2^ s^−1^, at 22±2°C; 8 h dark at 18±2°C). For experiments with leaf discs, 5-day-old seedlings were transferred to Jiffy-7^®^ peat pellets (Jiffy, Oslo, Norway) and grown under the same controlled conditions for 4–5 weeks. Leaves were excised using a 7 mm-diameter cork borer.

### Plant treatments

Treatments to evaluate gene expression were performed on 2-week-old seedlings grown under a long-day regime. Whole seedlings were collected at indicated time points, frozen in liquid nitrogen and stored at −80°C until processing. For SA treatments, seedlings were floated on a solution of 0.5 mM sodium salicylate (Sigma), 0.5x MS, 1% w/v sucrose liquid medium or 0.5× MS, 1% w/v sucrose liquid medium as control. Treatments with MV (Sigma) were performed with seedlings floated on a solution of 50 μM MV, 0.5× MS, 1% sucrose liquid medium or using 0.5× MS, 1% sucrose liquid medium as a control, for the indicated time points. MV stock aliquots (10 mM, dissolved in deionized water) were stored at −20°C in the dark and used within one month after the preparation. UV-B treatment was performed in a chamber equipped with two F8T5 UVP3400401 fluorescent tubes (λ = 306 nm) (USHIO, Cypress, CA) at a distance of 53 cm from the samples, irradiating with an intensity of 0.220 mW cm^−2^. As a control, a 2-mm- cellulose acetate polyester was used to filter UV-B radiation.

### Gene expression analysis

Total RNA was extracted with TRIzol® reagent (Invitrogen, Carlsbad, CA) following the manufacturer’s instructions. cDNA was synthesized from 2 μg of total RNA with the MMLV kit (Promega, Madison, WI). Semi-quantitative PCR was performed within the linear amplification phase using a recombinant Taq Polymerase (Invitrogen) amplifying a fragment of 323 base pairs (bps) of the coding sequence of *GSTU7* and a fragment of 294 bps of the *YLS8* housekeeping gene. Quantitative PCR was done using the Brilliant III Ultra-Fast SYBR® Green Master mix (Agilent Technologies, Santa Clara, CA) in an AriaMx (Agilent Technologies) or an MX3000P® instrument (Stratagene San Diego, CA). Expression levels of *GSTU7* were calculated as fold-change to the *CLATHRIN ADAPTOR COMPLEX* gene (*CLA)*, for SA and MV treatments, or to *YSL8* for UV-B treatments. Primer sequences are listed in Supplementary Table 1.

### Methyl viologen tolerance assay and ROS detection

MV tolerance assays were performed according to Laporte et al. (2012), with the following modifications. Surface sterilised seeds were stratified for 48 h in the dark at 4°C and then sown on MS medium solidified with 0.3% phytagel and supplemented with 0.1 μM MV. After one week, tolerance to MV was measured as survival rate that is the percentage of green seedlings/cotyledons compared to the total number of germinated seeds. The fresh weight (FW) from all germinated seedlings was measured after two weeks of growth.

To evaluate the mitigation of oxidative damage caused by MV, 2 μL of a 30 μM MV solution was placed as a single drop on the adaxial side of leaves of two-week-old seedlings grown on plates. The accumulation of H_2_O_2_ was measured after staining with DAB (Sigma) according to (Daudi et al., 2012). Leaves treated with MV were excised and placed in the dark in 6-well plates with 2 mL of staining solution. Samples were then vacuum-infiltrated for 5 min and incubated in orbital-shaking for 6 h or until staining was visible. Chlorophyll was removed by boiling in 80% ethanol and subsequent rinsing with distilled water. The stained leaves were mounted in 20% glycerol between two SDS-page glass plates and documented with an Epson V850 Pro scanner equipped with a dual lens system. Quantification of the staining intensity was performed using the FIJI software (Schindelin et al., 2012). Since DAB staining generates a dark precipitate, images were converted to 8 bit grayscale and the threshold was set on percentile and adjusted to indicate the darkest or less intense 3% of pixels in the image. Using the polygon selection, a region of interest (ROI) was selected around each leaf and saved with the ROI manager. The percentage of pixels below the threshold was measured over the total area of the ROI and expressed as “% DAB staining”.

### Cloning and plant transformation

For complementation of *gstu7* and transient expression of the GSTU7-GFP fusion protein, the constructs *UBQ10*_*pro*_*:GSTU7-V5* and *35S*_*pro*_*:GSTU7-GFP* were used, respectively. To obtain these constructs, a fragment of 681 bps corresponding to the full CDS of *GSTU7* (stop codon excluded) was amplified by PCR (see Supplementary Table 1 for primers). The fragment was purified using a NucleoSpin® kit (Macherey-Nagel, Düren, Germany) and cloned into the pENTR/SD/D-Topo vector (Invitrogen, Carlsbad, CA). The entry clone pEN-GSTU7 was recombined into the pB7m34gW destination vector (Karimi et al., 2005), containing the sequence for the ubiquitin 10 promoter (*UBQ10*_*pro*_) and V5 tag (*UBQ10*_*pro*_*:GSTU7-V5*). For subcellular localization, the CDS of *GSTU7* cloned in the pEN-GSTU7 was fused to the GFP fluorescent protein sequence by recombination with the pK7FWG2.0 binary vector (Karimi et al., 2002), generating the *35S*_*pro*_*:GSTU7-GFP* construct.

For protein purification, the plasmid pETG*-GSTU7-His* was generated. For this purpose, the CDS of *GSTU7* was amplified by PCR with primers carrying the *attB* recombination sites (see Supplementary Table 1); the fragment was then purified and cloned into the pDONR201 entry vector (Invitrogen) and recombined with the pETG10A vector, adding a C-terminal His_6_ tag to *GSTU7*. The entry vector, pDONR-*GSTU7*, was then recombined with the binary vector pAG415GPD-ccdB-HA (a gift from Susan Lindquist, Addgene plasmid # 14242) for *GSTU7* expression in *Saccharomyces cerevisiae*. All constructs were verified by sequencing.

For the transformation of *gstu7* mutant plants, the *UBQ10_pro_:GSTU7-V5* vector was transferred into the *Agrobacterium tumefaciens* strain C58C1 and stable Arabidopsis transgenic lines were generated by floral dip (Clough and Bent, 1998). Transformed seeds were surface-sterilized and selected on plates with 0.5x MS solid medium supplemented with 10 μg mL^−1^ ammonium glufosinate (Sigma).

To express the Grx1-roGFP2 redox sensor in the different genetic backgrounds, *gstu7* was crossed with a WT plant expressing Grx1-GFP2 in the cytosol, and the complemented line *gstu7/GSTU7* #5 was transformed by floral dip with a plasmid containing the *UBQ10_pro_*:Grx1-roGFP2 construct.

### Yeast complementation and growth

The *S. cerevisiae* mutant *gtt1::TRP1 gtt2::URA3* (Δ*gtt1/gtt2*), in the genetic background W303 (Choi et al., 1998; Collinson and Grant, 2003), was transformed either with the empty vector (ev) pAG415GPD-ccdB-HA or a version harboring *GSTU7* according to (Gietz and Woods, 2002). Cells were grown overnight at 30 C under constant agitation in 15 mL of synthetic SD medium without leucine (2% glucose, 0.67% yeast nitrogen base, 0.13% yeast synthetic drop-out supplement (Sigma), 0.004% L-histidin, 0.004% L-tryptophane, 0.002% L-uracil). Aliquots of the saturated cultures were incubated for 2 h in 15 mL of fresh medium until they reached exponential growth and then diluted with YPD media (20 g L^−1^ tryptone/peptone, 10 g L^−1^yeast extract) to a final OD_600_ = 0.1. Oxidative conditions were generated with H_2_O_2_ or CHP, added to the indicated final concentrations. Growth rates were measured as absorbance at 600 nm on a POLARstar plate reader (BMG Labtech, Ortenberg, Germany) in a final volume of 200 μL at 27.5°C.

### Protein extraction and immunodetection

Protein extracts were obtained from 2-week-old frozen seedlings by grinding the material in a microfuge tube with a plastic pestle. The resulting powder was suspended in lysis buffer (50 mM sodium phosphate buffer, pH 7.5–8.0, 150 mM NaCl, 0.2% IGEPAL and 5 mM EDTA) and supplemented with a protease inhibitor cocktail (Roche diagnostics). Samples were centrifuged at 2,350 *g* for 10 min at 4°C and the supernatant was collected. Protein concentration was determined by the Bradford method, using the BioRad Protein Assay (Bio-Rad Laboratories, Inc., CA, USA), and samples were stored at −80°C.

Immunoblot was performed according to (Seguel et al., 2018). Total protein samples (30 μg) were resolved in 12%-SDS-PAGE and transferred to ImmobilonTM PVDF membranes (Millipore, Burlington, MA) in a semi-dry electroblotting apparatus (Bio-Rad, Hercules, CA). Membranes were blocked overnight at 4°C in a solution of 5% (w/v) skimmed milk powder prepared in a PBS-T phosphate buffer (137 mM NaCl, 2.7 mM KCl, 10 mM Na2HPO4, 2 mM KH2PO4 and 0.1% (v/v) Tween-20). GSTU7-V5 fusion protein was detected with anti-V5 mouse monoclonal antibody (1:5,000 dilution, from Invitrogen #R96025) and a peroxidase-conjugated anti-mouse secondary antibody (1:10,000 dilution from KPL, Gaithersburg, MD #074-1806). GSTU7-V5 fusion was visualized with a 1:1 mix of SuperSignalTM West Pico and SuperSignalTM West Femto chemiluminescent substrates (Thermo Fisher Scientific, Waltham, MA).

### Purification of recombinant proteins and enzyme assays

A culture of *Escherichia coli* strain BL21 harboring pETG10A-GSTU7 was grown in LB medium supplemented with 100 μg mL^−1^ ampicillin at 37°C with constant shaking. Fresh cultures were prepared from saturated aliquots and grown until cells reached an OD_600_ of 0.6– 0.8. GSTU7-His expression was induced by incubating the cells for 24 h with isopropyl ß-D-1-thiogalactopyranoside (IPTG) in a final concentration of 0.5 mM and 200 μL of anti-foaming solution (Sigma), with constant shaking at 22°C. Cells were collected by centrifugation at 4,000 *g* for 15 min at 4°C and the pellet was suspended in a buffer (100 mM Tris-HCl, pH 8.0, 150 mM NaCl, 0.5 mM PMSF) supplemented with 5 mg L^−1^ DNAse I and 0.5 g/L lysozyme (Roche, Mannheim, Germany). Cells were disrupted by sonication (3×2 min, 40% power output, 50% duty cycle, in a SONOPULS Ultrasonic Homogenizer HD 2200, Bandelin, Berlin, Germany). The lysate was centrifuged at 19,000 *g* for 20 min at 4°C and the supernatant filtered through a sterile filter with 0.22 μm nominal size. The filtered fraction was loaded onto a Ni-NTA HisTrapTM column (GE Healthcare, Little Chalfont, UK) using a peristaltic pump at a rate of 1 mL min^−1^. Proteins were eluted from the column with a 10–200 mM imidazole gradient (100 mM Tris-HCl, pH 8.0, 200 mM NaCl) using an ÄKTA Prime Plus chromatography system (GE Healthcare, Chicago, IL). Fractions were collected and stored at 4°C and the presence of GSTU7-His protein was confirmed by analyzing the collected fractions by SDS-PAGE.

GPOX enzyme activity was measured using a GR-coupled protocol adapted from (Navrot et al., 2006) and expressed as the decrease in absorbance of NADPH at 340 nm in the presence or absence of a peroxide substrate. A final concentration of 10 μM recombinant GSTU7 protein, or no protein as the basal line, was incubated with 5 M yeast GR1, 400 μM NADPH and 500 μM GSH as the electron donor. The assays were carried out in a final volume of 100 μL in assay buffer (100 mM Tris-HCl, pH 7.4, 5 mM EDTA) where the substrates, H_2_O_2_ or CHP, were automatically injected to a final concentration of 200 μM. The absorbance was recorded at 340 nm in a POLARstar plate reader. The test thiol oxidase activity of GSTs, 10 μM of purified GSTU7 were incubated with 1 μM of purified roGFP2 and 4 mM of either H_2_O_2_ or CHP. Bovine GPX (Sigma) was used as a control. Oxidation of roGFP was followed on the plate reader exciting at 390 ± 5 nm and 480 ± 5 nm, and collecting the fluorescence from both channels at 520± 5 nm.

GST enzyme activity was measured as the conjugation of GSH to CDNB. The increase of GS-CDNB conjugate was followed as an increase in absorbance at 340 nm in a POLARstar plate reader. For preparing the stock solutions, 200 mM GSH was dissolved in assay buffer (100 mM Tris-HCl, pH 7.4, 5 mM EDTA) and 100 mM CDNB was dissolved in 96% ethanol. The assay mix was prepared by adding 100 μL of each stock solution to 9.8 mL of assay buffer before use. For the plate reader measurements, 195 μL of assay buffer were added to 5 μL of diluted recombinant GSTU7 protein and measured for 10 min after 10 s of orbital agitation at 200 rpm. All measurements were done in triplicates with 3 different batches of purified protein.

### Confocal microscopy

*Nicotiana benthamiana* plants were grown in soil for 3–4 weeks and leaves were transiently transformed by infiltration with *A. tumefaciens* as described by (Sparkes et al., 2006) using the bacterial strain AGL1 harboring the *35S_pro_:GSTU7-GFP* construct. Leaves were imaged 2 days later on a confocal laser scanning microscope (Zeiss LSM 780, connected to an Axio Observer.Z1; Carl Zeiss Microscopy, Jena, Germany) using a 40× lens (C-Apochromat 40X/1.2 W Korr). GFP and chlorophyll were excited at 488 nm and fluorescence emission was collected at 499–553 nm and 630–691 nm, respectively. Ratiometric imaging and analysis of WT and mutant lines harboring the Grx1-roGFP2 sensor were performed as described by (Attacha et al., 2017) utilizing a custom-written MATLAB script (Fricker, 2016).

### Monitoring of roGFP2 fluorescence in leaf discs

Fluorescence of the Grx1-roGFP2 sensor expressed in 4–5-week old WT, mutant and complemented lines was initially confirmed in a stereomicroscope (M165 FC, Leica, Wetzlar, Germany) equipped with a GFP filter (470 ± 40 nm excitation and 525 ± 50 nm emission) and documented with the attached camera (DFC425 C, Leica). Leaf discs were cut from positive fluorescent plants using a 7 mm cork borer and placed with the abaxial side up in the bottom of transparent Nunc^®^ 96-wells plates (Sigma).

For spectral measurements, leaf samples were excited in the range of 360–496 nm with a step width of 1 nm and the emission was collected at 520 nm using a CLARIOstar plate reader (BMG Labtech). Non-fluorescent samples of each genotype and treatment were used to subtract the basal background fluorescence.

Ratiometric measurements of Grx1-roGFP2 in leaf samples were performed in a POLARstar plate reader equipped with two excitation filters, 410 ± 5 nm for the peak of the oxidized roGFP and 480 ± 5 nm for the peak of the reduced roGFP; the emitted fluorescence was collected at 520 ± 5 nm. Samples were equally treated in 200 μL of final volume and the fully reduced and oxidized treatments were performed using a solution of 10 mM DTT or 5 mM DPS, respectively.

Leaf samples were treated with DTT to fully reduce the sensor, then rinsed twice with distilled water and treated with 200 μL of 100 μM of MV or distilled water as control. Fluorescence was measured every 210 s for over 15 h at a constant temperature of 25°C. To assess the dynamic range of the sensor, at the end of each experiment the probe was fully reduced and then fully oxidized by discarding the solution containing MV, and replacing it with 200 μL of 10 mM DTT for 5 min and then replacing it with 200 μL of 5 mM DPS. For every measurement, non-fluorescent samples of each genotype were treated and recorded under the same conditions. These values were later used to subtract autofluorescence. The experiment was repeated 6 times and, for each experiment, 6 leaf discs from independent plants were used for the measurements.

## Accession numbers

All the sequence information in this article is referred as in the GenBank/EMBL database under the following accession numbers. From *Arabidopsis thaliana*: AT2g29420 (*GSTU7*), AT1G17180 (*GSTU25*), AT5G06950 (*TGA2*), AT5G06960 (*TGA5*), AT5G06950 (*TGA6*), At5g46630 (*CLATHRIN ADAPTOR COMPLEX*), At5g08290 (*YLS8*), AT1g74710 (*ICS1/SID2*). From *Saccharomyces cerevisiae*: YIR038C (*GTT1/GST1*) and YLL060C (*GTT2/GST2*).

## Supplemental Data

**Supplemental Figure S1.** Unrooted phylogenetic tree of the GST protein family from Arabidopsis

**Supplemental Figure S2.** roGFP2 thiol oxidation assay.

**Supplemental Figure S3.** Subcellular localization and spectral properties of the Grx1-roGFP2 sensor in Arabidopsis leaves.

**Supplemental Figure S4.** Biological replicates for measurement of the glutathione redox state in response to MV in Arabidopsis leaves.

**Supplementary Table 1.** List of primers used in this work.

## Acknowledgements

We thank Dr. Chris Grant (University of Manchester) for providing the yeast strain Δ*gtt1*/*gtt2*, and Dr. Xin Li (University of British Columbia) for providing the *tga2-1 tga5-1 tga6-1* mutant line.

